# Survey on Prevalence of Canine Cutaneous Myiasis in Some Selected Kebeles of Dire Dawa City Administration

**DOI:** 10.1101/110502

**Authors:** Henok Abebe

## Abstract

A cross sectional study of canine cutaneous myiasis was conducted in five randomly selected kebeles of Dire Dawa Administrative council from December 2009 up to April 2010 to determine the prevalence of canine cutaneous myiasis and to assess factors that determine the occurrence of the disease specifically in dog. Active questionnaire survey among 60 households were used for which 384 dogs were sampled. From a total of 384 dogs, 162 (42.19%) were found harboring the disease cutaneous myiasis among this 120 (74.07%) were infested with the 3^rd^ and 2^nd^ instar larvae of *Cordylobiaantropophaga.* whereas the remaining 42(25.93%) observed dogs were found infested with cutaneous myiasis. The larvae were identified in Dire Dawa regional diagnostic veterinary parasitology laboratory. Analysis of active questionnaire survey showed that there is no statistically significance difference in the prevalence of disease among different breeds and sexes (P >0.05). In this study, an overall prevalence rate of 162 (42.19%) was found with a statistically significant association among different age groups, housing system and living area (kebele) (P<0.05). Higher prevalence was recorded at 02 kebele (Sabian area) 59 (54.65%), 03 Kebele (Depo and number-one), 43(44.33%), 04 Keble (Gende kore and Greek camp) 33(37.50%), Addis Ketema. 27(51.92%) and05 Keeble (Dechatu) 0(0%). There was 121(49.59%) confined dogs and 41(29.29%) were stray dogs which let out without any control, and puppies of age less than 6 month old (71.56 %), and dogs of age range between 6 months and 18months (79.03%) while those of greater than 18 months (16.43%), were least affected.

## 1. INTRODUCTION

Myiasis is infestation of human and vertebrates with the larval stage of dipterous flies. It has been known to occur at a various site of the body such as eye, intestine, mouth, nose, urogenital tracts, and brain where they survive by feeding on the living or dead tissues, ingested food or body fluids **[2, 20]**

Myiasis can occur in all sorts of variations depending on fly species and where the larvae are located. Some flies may lay egg in open wound, others’ larvae may invade unbroken skin or enter the body through the nose or ear and still other may be swallowed if eggs are deposited on the lip or on food **[7]**.

*Cordylobia* species *Cordylobiaantropophaga* is a large robust, brownish-yellowish fly (i.e.) “tumbu fly” of Africa which causes a boil like (funicular) type of myiasis particularly of human and dogs. Eggs are deposited on dry, shaded ground, specially; if contaminated with animal’s faces and urine which then attracts flies to lay egg; those eggs hatch after 1-3 days and remain just under soil surface until activated by host’s body heat. Then they emerge and actively penetrate normal skin of the host and burrow in a furuncle **[4]**

The larva remains in the subcutaneous tissue until it reaches maturity feeding on liquid protein [3] without migrating to deeper structures for a set time, it molts to second stage larvae which feed for a further period before moulting to third stage larvae. The larva in this stage feed voraciously for a set of time and then leaves the food sources in search of a safe and dry place in which to pupate away from the animal. The pupa forms an outer coat and metamorphosis occurs after a few days, the adult fly emerges and the empty puparium is left behind **[9]**

Opening to the skin allows it to breathe and eliminate larval excretions **[16]** the larva possesses two spiracles that give origin to two tracheas **[18]**, and these are located in the posterior portion of the larva, close to the skin **[10]**

Cases of dog myiasis are frequent especially in underdeveloped countries. It also has been described in human who reported trips to tropical Africa or who live there, where the disease is referred as *“tumbu”* **[11]**

Dog and small rodents are particularly important reservoirs for the parasite, Humans infected only accidentally. Only few cases have been reported in the literature and the majority of them are due to the human botfly *Dermatobiahominis* **[17]**. To the best of our knowledge, only one case of breast myiasis due to *Cordylobiaantropophaga* has been reported in the literature, a 70 years old women from South East Nigeria where referred to the surgery clinic of Albi State University Hospital with a week history of “multiple itchy, painful boil” of right breast **[3]**.

Elderly people who are ill or debilitated; they are especially seen in the tropics and the Third World countries. Although all age groups may be affected, the damage caused to the infants is more serious and may even be fatal **[13]**

Typical clinical sign observed in canine cutaneous myiasis include numerous erythematous, frunculoid skin lesions which oozes serous fluid. Each nodular lesion contains one larva although multiple lesions may occurrarely more than four or five lesions application of a pressure to such lesion may lead to the larva (e) with liquefied hemorrhagic or purulent tissue **[15]**

Symptoms of myiasis in human case are usually observed during the 1^st^ week of clinical evaluation if there are multiple sites of infestation, hosts may develop enlarged lymph nodes and fever. If lesions are numerous, they may coalesce forming large plaques with a serious thin, yellow discharge **[6]**

Diagnosis is mainly clinical. The main diagnostic features are recent travel to an endemic area, one or more nonhealing lesions on the skin, symptoms of pruritus, movement under skin or pain. Other features include serous or serosanguinous discharge from a central punctum and small, white thread-like structure protruding from the lesion. Definitive diagnosis of canine myiasis is usually by the extraction of the larvae and laboratory identification of the larvae. **[8]**

The control and prevention of cutaneous myiasis in animals mainly depends on application of strict hygienic control and immediate treatment of all infested dogs with approved insecticides. Animals should be quarantined until the wound clearly healed and it is also important to inform local residents especially dog owners the need to remove the rubbish around home. Regular and proper disposal of carcasses, feces, and decomposing materials, all of which serve as breeding sites for the fly and bedding on which flies may deposit their egg. **[1]**

Myiasis is considered to be endemic in tropical Africa. Funicular myiasis because of *CORDYLOBIAANTROPOPHAGA* infestation has been endemic in the West African sub region for more than 135 years. The high prevalence of the disease in the continent is probably because many African countries have not yet started the control and eradication schemes; however, few countries have carried out extensive survey on dog’s *cutaneous myiasis.* For instance prevalent rate of 58.95% in females and 41.05% in male with intensity level 517(54%) and 440(46%) larvae extracted with statistically significance difference (p<0.05) in Nigeria was documented (The prevalence rate of 68.9% in young dogs of age less than 5month were diagnosed with *myiasis* **[11]**.

In Ethiopia despite its potential public health significance of canine myiasis, there is no published report of the actual fly species causing this condition in the study area. Therefore, the objectives of this study are to estimate the prevalence of canine cutaneous myiasis in Dire Dawa Administrative council state and to assess the apparent densities of the canine cutaneous myiasis and identification of larvae and factors that predisposes dogs to cutaneous myiasis.

## 2. MATERIALs AND METHODES

### 2.1. Description of Study Area

The study was conducted in5 randomly selected kebeles of Dire Dawa administrative council from December 2009 up to April 2010. Dire Dawa is situated about 518 Km east of Addis Ababa, between 09’ 28’’ to 09’49’’N latitude and 41’38’’ to 42’ 19’’E longitude. The land is situated in the altitude range of 950-2250m a. s. l. The rainfall pattern is bimodal with the highest rainfall in July and August with average of 700 mm to 900 mm. The monthly mean maximum temperature range up to 28.1℃ recorded in May. Total area of the administrative council is about 1288.02 kilo meter square and the total livestock population is estimated to be about 37,129 heads of cattle, 153,778 heads of small ruminants, 7,513 camels, 10,779 equines, 25,301 chicken and 1,225 bee hives **[14]**

### 2.2. Study Population

The study animal comprises local, exotic and hybrid dogs of different age group of both sex in a different management system, were indigenous breeds of dog presented from a wide range of owners in selected kebeles to ensure random selection of animal.

### 2.3. Study Design

A cross sectional study was carried out to estimate the prevalence of canine cutaneous myiasis and to assess the factor that facilitates the occurrence of the disease. The study is limited mainly on the dog in the study area based on laboratory examination and active questionnaire survey.

#### 2.3.1. Sampling Methods

The sample size determination was preformed according to the method previously provided byThrusfield (2005). The following formula was used to calculate the number of animals to be sampled.

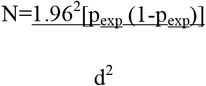

An expected prevalence 50% was used and considering 5% precision and 95% CI gave as 384-sample size. Therefore, a total of 384 animals were sampled and examined.

Where:

N=required sample size
p=expected prevalence
d^2^=despaired absolute precision

By taking, an expected prevalence of 50% at 95% confidence interval and 5% desired absolute precision since there is no previous data on prevalence of the disease 384 dog population was determined.

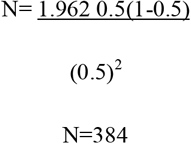

A simple random sampling technique was used to collect data on the presence of the disease and to estimate its prevalence.

#### 2.3.2. Questionnaire Survey

Active questionnaire was conducted on five randomly selected kebeles as a preliminary investigation to estimate the prevalence of the disease. A total of 100 households were interviewed and less aggressive infested dogs were secured and larvae samples was collected whereas the rest investigated by asking the owner regarding housing, breed, sex, and age of the dog, knowledge of the disease and if there is case of myiasis in humans (Annex 2).

**Annex 1:**
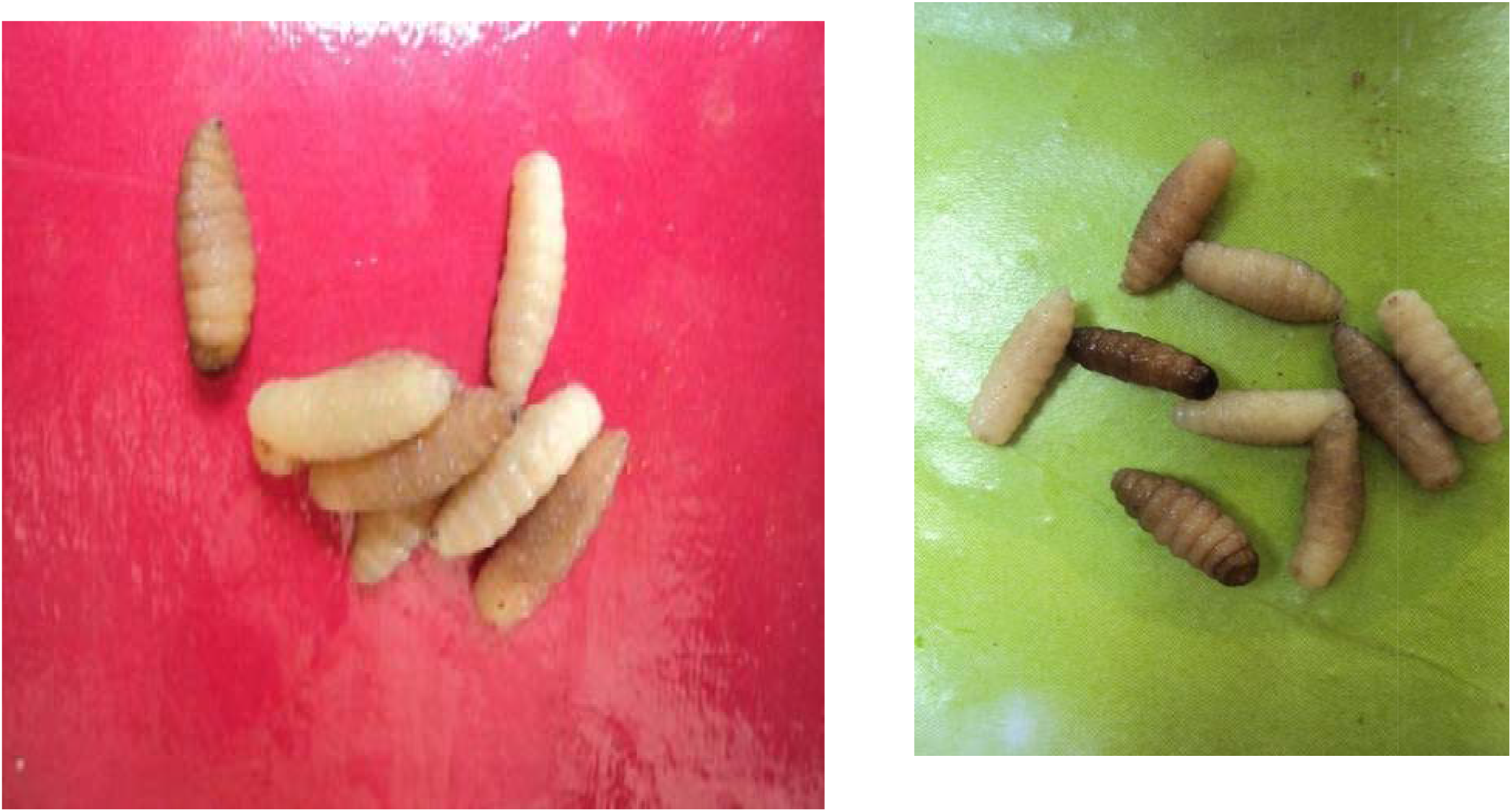
Image of a *Cordailobia antropophaga* larvae sample from dog.

#### 2.3.3. Laboratory Examination

Laboratory examinations were carried out in Dire Dawa regional diagnostic veterinary laboratory. Out of the total population, 122 larvae samples was collected from myiasis-infested dogs of various breeds, different age groups of both sexes. The larvae sample collected from each dog were put into 10 ml bottle containing 10% formal slain solution as a preservative and information’s such as breed, age and sex of dogs, number of larvae harvested and addresses were written and taken to the regional parasitology laboratory for identification. Identification was carried out using posterior spiracle plate method according to **[15]**

In brief each larvae was boiled in 10% KHO for 5 minutes and posterior spiracle cut to the depth of about 1 mm, excess tissue around the plate was treasured out before being dehydrated in varying concentration of alcohol (70%-100%), xylene clinging took place before mounting and observation under the dissecting microscope **[12]**

### 2.4. Data Management and Analysis

The data was entered into a Microsoft Excel spread sheet and statistical analysis was carried out using STAT 7.0 version software. Categorical variables (breed, sex, housing, kebeles and age) were expressed in percentage. The prevalence rate of cutaneous myiasis of dogs calculated by dividing total number of dogs harboring the larvae by the total number of dogs examined. For data analysis 95% confidence interval and 5% level of precision was considered. Percentage (P. value) was considered to measure occurrence of association between variables and occurrence of the disease.

## 3. RESULTS

Out of 384 dogs (262 male) and (122 female) examined for cutaneous myiasis larvae, 162 (42.19%) were found to be positive for larvae of *Cordylobiaantropophaga.* From the finding the result of the present study revealed that there is statistically significance difference among dogs of different management or housing system (P< 0.05) (table 1).

**Table 1:** Canine cutaneous myiasis in different housing system.

| *Housing system* | *Number examined* | *Prevalence* | *X<sup>2</sup>* | *P. Value* |
| --- | --- | --- | --- | --- |
| Indoor dogs | 243 | 121 (49.59%) | 15.037 | 0.000 |
| Stray dogs | 140 | 41 (29.29%) |  |  |
| **Total** | **384** | **162 (42.19%)** |  |  |

The overall prevalence of canine cutaneous myiasis in the present study was 42.19 % (162/384), from which the respective canine cutaneous myiasis prevalence in 02, 05 04, 01, and 06 kebele includes (54.92%), (51.92%), (44.33%), (37.50%) and (0%) in descending order of prevalence. Table 2

**Table 2:** Prevalence of canine myiasis in five randomly selected kebeles of Dire Dawa.

| *Kebele* | *Examined animal* | *Positive* | *X<sup>2</sup>* | *P. Value* |
| --- | --- | --- | --- | --- |
| 02 | 108 | 59 (54.92%) | 38.31 | 0.000 |
| 01 | 88 | 33 (37.50%) |  |  |
| 04 | 97 | 43 (44.33%) |  |  |
| 05 | 52 | 27 (51.92%) |  |  |
| 06 | 39 | 0 (0%) |  |  |
| **Total** | **384** | **162 (42.19%)** |  |  |

The overall prevalence of canine cutaneous myiasis in male and female sex groups was 39.31% and 48.36%, respectively. The difference in overall prevalence between male and female dogs is higher in female than male dogs was statistically significant (p<0.05) (Table 3).

**Table 3:** Sex distribution of canine cutaneous myiasis infestation of dogs in Dire Dawa.

| *Sex* | *No. examined* | *Positive* | *X<sup>2</sup>* | *P. Value* |
| --- | --- | --- | --- | --- |
| Male | 262 | 103 (39.31%) | 2.7938 | 0.095 |
| Female | 122 | 59 (48.36%) |  |  |
| **Total** | **384** | **162 (42.19%)** |  |  |

Prevalence of canine cutaneous myiasis between local (44.03%), hybrid (30.88%) and exotic breed (41.18%) were analyzed and there is no significant difference between the breeds (p>0.05). (Table 4).

**Table 4:** Breed distribution of canine cutaneous myiasis in dog examined in Dire Dawa.

| *Breed* | *No. of animal* | *Positive* | *X<sup>2</sup>* | *P. value* |
| --- | --- | --- | --- | --- |
| Local | 299 | 134 (44.03%) | 4.4175 | 0.11 |
| Hybrid | 68 | 21 (30.88%) |  |  |
| Exotic | 17 | 7 (41.18%) |  |  |
| **Total** | **384** | **162 (42.19%)** |  |  |

In this study the overall prevalence of canine cutaneous myiasis was analyzed and higher in young, puppy and adults was 79.03%, 71.56% and 16.43% respectively. The difference was statistically significant (p<0.05) (Table 5).

**Table 5:** Prevalence of canine cutaneous myiasis based on age groups.

| *Age Category* | *No. examined* | *Positive* | *X<sup>2</sup>* | *P. Value* |
| --- | --- | --- | --- | --- |
| Less than 6 month | 109 | 78 (71.56%) | 130.99 | 0.000 |
| 6-18 month old | 62 | 49 (79.03%) |  |  |
| Greater than 18 month | 213 | 35 (16.43%) |  |  |
| **Total** | **384** | **162 (42.19%)** |  |  |

## 4. DISCUSSION

The overall prevalence of canine cutaneous myiasis in the current study is found to be (42.19%). There is no published and recorded research works in Ethiopia even to compare with present work, yet the result was found in fair agreement with report **[11]**, 77.50 % in Nigeria where 54% female and 46% male dogs were found positive. **[4, 13]** also 68.9% reported in the same country where 58.95% puppies with intensity level of 4.62 % larvae.

However, in the present study there is no statistical significant difference between sexes and between deferent breeds of dogs and it seems to contradict with established facts. However, it is difficult to make firm conclusion, as the number of study animals is small in proportion as compared to the factors. The prevalence would have been higher in immature puppies and young female dogs, as **[4]** reported that the infestation was high in young dogs because of their thin, soft, skin, which has been found suitable for larval development.

In the present study, the occurrence of the disease in different age categories was assessed. In the adult age, the prevalence was found low as the result of low exposure to the ground where the egg of the *“tumbu fly”* rest but conversely on the younger and puppies the prevalence of cutaneous myiasis was higher due to routine contact with ground as well as much more heat dissipation that may attract the larvae. The prevalence of 71.56% was recorded in puppies of age less than six month old whilst 79.03% in the dogs of age ranging between six and eighteen months and 16.43% in dogs of age greater than eighteen month old. The difference were statistically significant between age groups (<0.05) which is in agreement with the result reported in the study in Britain on pet myiasis **[5]**

As observed in the present study, young dogs were usually maintained in the house and they do not go here and there. This decreases the chance of defecation and urination at the same place repeatedly that may contaminate the area and creates suitable area for the fly to lay eggs and the larvae had easy accessible to host and can penetrate the soft tissues of young and puppies. This is also supported by reports of **[15]**

The finding of the present study revealed that there is statistically significance difference (P <0.05) among dogs of different management or housing system (table 1).

## 5. CONCLUSIONS AND RECOMMENDATIONS

The result of this study revealed that there is a wide spread of canine cutaneous myiasis in the study area with an overall prevalence rate of 42.19%. Such findings of moderately high prevalence indicate that the occurrence of a natural infection. This study also indicates that there is a tradition of keeping dogs under a different management system for several intended use were as less attentions are given to the predisposing factors and the public health significance of the disease. This may be due to the lack of awareness about the etiology and predisposing factors canine cutaneous myiasis since larvae of *Cordylobiaantropophaga “tumbu fly”* of Africa in which the health of dogs is endangered by the infestation. Children and women have been found contracting myiasis as they carry out most activities related with pet animals.

In line with the above conclusion, the following recommendation are forwarded

✓ Further epidemiological studies and identification of the fly species responsible for myiasis shall be carried out.
✓ There should be awareness creation about the public health importance of the myiasis.
✓ There should be coordination among the veterinarians, public health officers and the concerned municipality officials for other further investigation of the case and make decisions on the way forward.

